## Supplemental text for "Transcription factor competition facilitates self-sustained oscillations in single gene genetic circuits"

(Dated: June 14, 2022)

### S1. THE TROUBLE WITH HILL FUNCTIONS

In fig. S1 we show how the Hill function is able to approximate the occupancy $\theta$ of a binding site in the presence and absence of a number of competitor sites. In the absence of

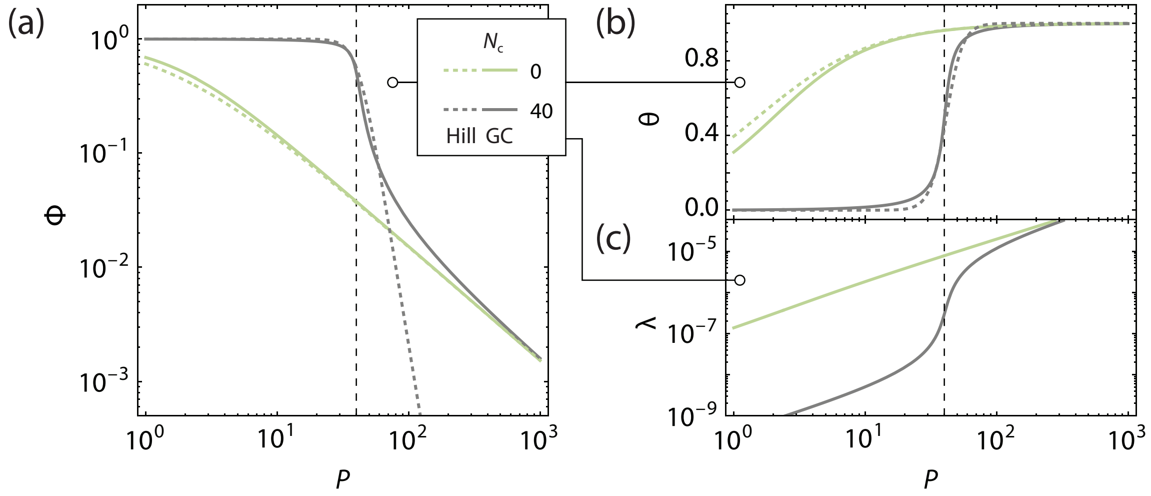


FIG. S1. **The trouble with Hill functions. a** Fold-change of a gene regulated by a simple repression scenario as a function of repressor copy number $P$, in the presence and absence of competitor sites is captured accurately by the grand canonical formalism. Using Hill functions, the Langmuir-behaviour in the absence of competitors can be accurately described. However, the Hill equation can not capture the full behaviour of the titration effect caused by competitor sites, even when fitting both the Hill exponent and the binding free energy. The small mismatch between the grand canonical and Hill curves at low transition factor copy number is a known effect of the differences in statistical mechanical ensemble in which these curves were calculated[1]. **b** The occupancy $\theta$ of repressors to their binding sites is captured well by the Hill equation. **c** The dramatic switching behaviour is a consequence of the lowered transcription factor fugacity at concentrations lower than the number of competitor sites. The binding free energy of repressors was kept at $-15\,k_\mathrm{B}T$ and for competitor sites at $-18\,k_\mathrm{B}T$. The number of non-specific sites was kept at $5 \times 10^6$. The Hill function in the presence of competitors was given an effective binding energy of $-11.3\,k_\mathrm{B}T$ and Hill exponent of 7. The model used to calculate these curves is presented in Section S2


*

†

competitor sites, transcription factor binding follows the Langmuir isotherm. In the presence of 40 competitor sites, the sharp switching behaviour of the occupancy can be captured by the Hill function, yet both the Hill exponent and the binding free energy have now become fitting parameters of obscure physical relevance, and still the resulting function does not capture the full fold-change behaviour.

### S2. FINDING THE TRANSCRIPTION FACTOR FUGACITY

In general, transcription factors can bind to their specific binding sites related to $N \geq 1$ copies of the gene of interest, to the reservoir of $N_{\text{ns}} \gg 1$ non-specific binding sites, or to a set of additional reservoirs $\{k\}$, each with $N_k$ binding sites, which can be binding sites related to competitor genes. Individual molecules can transfer between reservoirs, while the total number of molecules in the cell is conserved. When needed, a reservoir for free transcription factors can be included. However, as mentioned before, the fraction of transcription factors unbound to DNA is generally negligible, hence our choice is to not to include a reservoir for free transcription factors in solution. The fugacities $\lambda_1, \lambda_2, \ldots$ are set by the constraint that mass is conserved inside the cell, and can be found by setting up a mass balance that contains all relevant reservoirs. In general, that mass balance equation takes the form

$$P_i = N\theta_i + N^{(\text{ns})}\theta_i^{(\text{ns})} + \sum_{\{k\}} N^{(\text{k})}\theta_i^{(\text{k})}. \tag{1}$$

with $\theta_i, \theta_i^{(\text{ns})}$ and $\theta_i^{(k)}$ the occupational fractions of transcription factor $i$ to specific sites, non-specific sites and $k$-sites respectively.

#### A. Simple repression

We work through the examples given in the main paper to show how this mass balance is set up and solved, starting with the architecture known as 'simple repression' in the absence of competitor genes. Here, the occupational fractions of the repressor molecule to its specific site ($\theta_i$) and to non-specific sites ($\theta_i^{\text{ns}}$) are given by

$$\begin{cases} \theta_i = \dfrac{\lambda_i x_i}{1+\lambda_i x_i + \lambda_1 x_1} \simeq \dfrac{\lambda_i x_i}{1+\lambda_i x_i}, & (\lambda_1 x_1 \ll 1) \\ \theta_i^{(\text{ns})} = \dfrac{\lambda_i}{1+\lambda_i} \simeq \lambda_i. \end{cases} \tag{2}$$

Here we have introduced the shorthand $x_i = \exp(-\beta\epsilon_i)$. In the situation that $\lambda_1 x_1 \ll 1$, this RNAP-related term can be neglected and the occupational fraction of the repressor molecule loses its coupling to the RNAP fugacity. It turns out this situation is rather general, and is commonly referred to as the 'weak promoter limit'[2]. Furthermore, we have set the reference chemical potential of the transcription factors to equal the effective binding free energy of the non-specific reservoir $\epsilon_{\text{eff}}^{(\text{ns})}$. One is free to choose this reference as one chooses, and this choice simplifies the notation.

The mass-balance equation in this architecture now takes the form

$$P_i = N\frac{\lambda_i x_i}{1+\lambda_i x_i} + N^{(\text{ns})}\lambda_i, \tag{3}$$

which is a quadratic equation in $\lambda_i$ which has two solutions

$$\begin{aligned} \lambda_i = & \frac{(P_i - N)x_i - N^{(\text{ns})}}{2N^{(\text{ns})}x_i} \\ & \pm \frac{\sqrt{(N^{(\text{ns})})^2 + 2NN^{(\text{ns})}x_i + 2N^{(\text{ns})}P_i x_i + N^2 x_i^2 - 2NP - ix_i^2 + P_i^2 x_i^2}}{2N^{(\text{ns})}x_i}. \end{aligned} \tag{4}$$

Only the positive root gives a physical solution, so we discard the negative root. We can see that this equation describes the fugacity for any number $N$ gene copies, since their presence was explicitly included in the mass balance equation. Its effect is a sharp shift in transcription factor fugacity when the transcription factor copy number roughly equals the gene copy number.

#### B. Competitor sites

When an additional set of $N^{(\text{c})}$ competitor genes is present, competing for the same pool of transcription factors, an additional reservoir is added to the mass balance equation. A corresponding transcription factor occupancy $\theta^{(\text{c})}$ is calculated from the competitor gene's particular regulatory architecture. For simplicity, we assume again the same 'simple repression' architecture, although the reservoir can be set up to accomodate additional complexity.

The transcription factor occupancy to the competitor reservoir is given by

$$\theta_i^{(c)} \simeq \frac{\lambda_i x_i^{(c)}}{1 + \lambda_i x_i^{(c)}}, \tag{5}$$

where $x_i^{(c)} = \exp(-\beta\epsilon_i^{(c)})$ with $\epsilon_i^{(c)}$ the binding free energy of transcription factor $i$ to the competitor gene. Including this reservoir in the mass balance equation,

$$P_i = N\frac{\lambda_i x_i}{1 + \lambda_i x_i} + N^{(ns)}\lambda_i + N^{(c)}\frac{\lambda_i x_i^{(c)}}{1 + \lambda_i x_i^{(c)}}, \tag{6}$$

leads to a cubic equation in $\lambda_i$ of the form $a\lambda_i^3 + b\lambda_i^2 + c\lambda_i - P_i = 0$, with the coefficients

$$\begin{cases} a = x_i^{(c)} x_i N^{(ns)} \\ b = (x_i^{(c)} + x_i)N^{(ns)} + x_i^{(c)} x_i (N + N^{(c)} - P_i) \\ c = N^{(ns)} + x_i^{(c)}(N^{(c)} - P_i) + x_i(N - P_i). \end{cases} \tag{7}$$

This cubic equation has three solutions, of which only one is both positive and real,

$$\lambda_i = \Delta_+ + \Delta_- - \frac{b}{3a}, \tag{8}$$

where, $\Delta_\pm = (C_2 \pm \sqrt{C_1^3 + C_2^2})^{1/3}$ with $C_1 = (c/3a) - (b/3a)^2$ and $C_2 = (bc/6a^2) + (P_i/2a) - (b/3a)^3$. The solution allows one to vary both the copy number of competitor genes, as well as the copy number of the gene of interest. Including more reservoirs increases the order of the polynomial to solve, and for more complex situations, one may have to resort to numerical solutions to the mass balance equation. Similarly, the presence of competitor genes causes a sharp transition in the transcription factor fugacity that occurs when the number of transcription factors roughly equals the number of competitor genes. This shift is called the titration effect[3] and is also visible in fig. S1.

As shown in the main paper, this solution also works for competitive inhibitors of the transcription factors, as such agents would also include an additional reservoir to the mass balance equation. If the transcription factor completely detaches from the DNA while bound to the competitive inhibitor, then this solution is exact. However, as is the case for IPTG binding to LacI, the transcription factor may remain bound to the non-specific reservoir, essentially shrinking the effective size of the non-specific reservoir. Provided the number of transcription factors or inducers is far smaller than the number of basepairs, this effect can safely be neglected.

#### C. Transcription factor looping

In the transcription factor looping scenario described in the main paper, the nonlinearity in the fold-change function comes from the architecture, which changes the form of the equation describing the occupancy of the transcription factor binding site. In the case of the looping scenario, the transcription factor has two binding heads that can bind to two independent binding sites near the promoter sequence of the gene. In general, only one of these binding sites prohibits binding of RNAP, and this site is referred to as the main (m) operator site, with the other referred to as auxiliary (a). This architecture allows additional configurational states as is described in e.g. [4, 5], and occurs in e.g., the well studied *lac* operon [6]. The grand canonical partition sum of a gene regulated by the looping architecture is given by

$$\Xi = 1 + \lambda_1 x_1 + \lambda_2 \left(x_2^{\mathrm{m}} + x_2^{\mathrm{a}} + x_2^{\mathrm{m}} x_2^{\mathrm{a}} e^{-\beta F_{\mathrm{L}}}\right) + \lambda_1 \lambda_2 x_1 x_2^{\mathrm{a}} + \lambda_2^2 x_2^{\mathrm{m}} x_2^{\mathrm{a}}, \tag{9}$$

where as before we have $\lambda_{1,2}$ the fugacities of RNAP and the transcription factor respectively, with $x_i = \exp(-\beta\epsilon_i)$ the Boltzmann exponent of the binding free energy of transcription factor $i$. We have explicitly labelled the binding free energy of the transcription factor to the main transcription factor binding site (m) and to the auxiliary binding site (a), which need not *a priori* be equal. The architecture allows simultaneous binding of RNAP and transcription factor, provided the transcription factor binds to the auxiliary site, and as such, the configurational state where both are bound is taken into account in the grand canonical partition sum. Finally, the simultaneous binding of both main and auxiliary sites by the same transcription factor is possible, but requires a physical DNA-transcription factor loop which costs a certain free energy penalty $F_{\mathrm{L}}$. This free energy penalty represents the reduced conformational degrees of freedom that the DNA can adopt while in this configuration, and depends on the length of the DNA-transcription factor loop [7].

The transcription factor occupancy $\theta_2$ of this gene follows by taking the appropriate derivative of the grand canonical partition sum with respect to the transcription factor fugacity, $\partial \log(\Xi)/\partial \log(\lambda_2)$, and is given by

$$\theta_2 = \frac{\lambda_2 \left(x_2^{\mathrm{m}} + x_2^{\mathrm{a}} + x_2^{\mathrm{m}} x_2^{\mathrm{a}} e^{-\beta F_{\mathrm{L}}}\right) + \lambda_1 \lambda_2 x_1 x_2^{\mathrm{a}} + 2\lambda_2^2 x_2^{\mathrm{m}} x_2^{\mathrm{a}}}{1 + \lambda_1 x_1 + \lambda_2 \left(x_2^{\mathrm{m}} + x_2^{\mathrm{a}} + x_2^{\mathrm{m}} x_2^{\mathrm{a}} e^{-\beta F_{\mathrm{L}}}\right) + \lambda_1 \lambda_2 x_1 x_2^{\mathrm{a}} + \lambda_2^2 x_2^{\mathrm{m}} x_2^{\mathrm{a}}}. \tag{10}$$

Without loss of generality, we can replace the subscript 2 by the generic transcription factor subscript $i$ here. First, we remark here that the occupancy is coupled to the fugacity of RNAP. In rather general situation that $\lambda_1 x_1 \ll 1$ — the weak promoter limit — this coupling is resolved and the transcription factor occupancy tends to

$$\theta_i \simeq \frac{\lambda_i \left(x_i^{\mathrm{m}} + x_i^{\mathrm{a}} + x_i^{\mathrm{m}} x_i^{\mathrm{a}} e^{-\beta F_{\mathrm{L}}}\right) + 2\lambda_i^2 x_i^{\mathrm{m}} x_i^{\mathrm{a}}}{1 + \lambda_i \left(x_i^{\mathrm{m}} + x_i^{\mathrm{a}} + x_i^{\mathrm{m}} x_i^{\mathrm{a}} e^{-\beta F_{\mathrm{L}}}\right) + \lambda_i^2 x_i^{\mathrm{m}} x_i^{\mathrm{a}}}. \qquad (\lambda_1 x_1 \ll 1) \tag{11}$$

Second, we remark that at high transcription factor fugacities, the occupancy tends towards 2, reflecting a situation where both the main and the auxiliary operator site are saturated with transcription factor.

In the absence of any competitor sites, the transcription factor fugacity can now be found by setting up the usual mass balance equation

$$P_i = N \frac{\lambda_i \left(x_i^{\mathrm{m}} + x_i^{\mathrm{a}} + x_i^{\mathrm{m}} x_i^{\mathrm{a}} e^{-\beta F_{\mathrm{L}}}\right) + 2\lambda_i^2 x_i^{\mathrm{m}} x_i^{\mathrm{a}}}{1 + \lambda_i \left(x_i^{\mathrm{m}} + x_i^{\mathrm{a}} + x_i^{\mathrm{m}} x_i^{\mathrm{a}} e^{-\beta F_{\mathrm{L}}}\right) + \lambda_i^2 x_i^{\mathrm{m}} x_i^{\mathrm{a}}} + N^{(\mathrm{ns})} \lambda_i, \tag{12}$$

which leads to a cubic equation of the form $a\lambda_i^3 + b\lambda_i^2 + c\lambda_i - P_i$, where the coefficients are given by

$$\begin{cases} a = x_i^{(\mathrm{m})} x_i^{(\mathrm{a})} N^{(\mathrm{ns})} \\ b = \left(x_i^{\mathrm{m}} + x_i^{\mathrm{a}} + x_i^{\mathrm{m}} x_i^{\mathrm{a}} e^{-\beta F_{\mathrm{L}}}\right) N^{(\mathrm{ns})} + x_i^{\mathrm{m}} x_i^{\mathrm{a}} \left(2N - P_i\right) \\ c = N^{(\mathrm{ns})} + \left(x_i^{\mathrm{m}} + x_i^{\mathrm{a}} + x_i^{\mathrm{m}} x_i^{\mathrm{a}} e^{-\beta F_{\mathrm{L}}}\right) \left(N - P_i\right). \end{cases} \tag{13}$$

This cubic equation again has three solutions, of which only one is both positive and real,

$$\lambda_i = \Delta_+ + \Delta_- - \frac{b}{3a}, \tag{14}$$

where, $\Delta_\pm = (C_2 \pm \sqrt{C_1^3 + C_2^2})^{1/3}$ with $C_1 = (c/3a) - (b/3a)^2$ and $C_2 = (bc/6a^2) + (P_i/2a) - (b/3a)^3$.

### S3. NUMERICAL INTEGRATION OF DDES AND CALCULATION OF THE STABILITY CONTOURS

A Mathematica 12[8] notebook is included as supplementary material to this paper, in which fold-change functions, analytical solutions to the transcription factor fugacity $\lambda_i$ and numerical integrations of the rate equations in the main paper can be calculated. The notebook is able to reproduce the figures from the main paper.

All numerical integrations of DDEs are performed using the method of steps implemented in the Mathematica function `ParametricNDSolve`.

The notebook also allows calculation of the stability contour and the bifurcation diagram in several experimental parameters. The calculation of the slope of the fold-change function $\Phi_i$ in the stationary point was calculated by first generating analytical expressions for the derivative of $\Phi_i$ as a function of the transcription factor copy number $P_i$ using Mathematica's symbolic maths evaluation. The stationary point in any circuit with only one protein type is always found as the root of the equation $P_i = \Phi_i(P_i)$ as the crossover point of the two nullclines.

### S4. MULTIPLE DELAY TIMES

The time delay $\tau$ was explicitly introduced in the process of transcription, so that the introduction of newly synthesised mRNA depends on the fold-change at the time of transcription-initiation. This mRNA is immediately available for translation. In reality both transcription and translation take a finite amount of time and it would therefore make sense to introduce an explicit delay in both differential equations. It turns out that this does not affect the stability of the gene circuit. The only factor of importance is the sum of all the delay times in the feedback loop. Dividing the delay over the transcription and translation process will only cause a phase shift in the concentration of mRNA and protein. To see why this is we write down the rate equations in the case the explicit delay is present only in the translation step. In that case, we have

$$\begin{cases} \dfrac{\mathrm{d}\mathbf{m}(t)}{\mathrm{d}t} = -\Gamma_{\mathrm{M}}\mathbf{m}(t) + \Gamma_{\mathrm{M}}\,\boldsymbol{\Phi}(\mathbf{p}(t)) \\ \dfrac{\mathrm{d}\mathbf{p}(t)}{\mathrm{d}t} = -\Gamma_{\mathrm{P}}\mathbf{p}(t) + \Gamma_{\mathrm{P}}\mathbf{m}(t-\tau). \end{cases} \tag{15}$$

If we now introduce $\mathbf{m}'(t) \equiv \mathbf{m}(t-\tau)$ as the normalised concentration of mRNA shifted in time by $\tau$, and substitute $\mathbf{m}'$ for $\mathbf{m}$ in eq. (15), we get

$$\begin{cases} \dfrac{\mathrm{d}\mathbf{m}'(t)}{\mathrm{d}t+\tau} = -\Gamma_{\mathrm{M}}\mathbf{m}'(t+\tau) + \Gamma_{\mathrm{M}}\,\boldsymbol{\Phi}(\mathbf{p}(t)) \\ \dfrac{\mathrm{d}\mathbf{p}(t)}{\mathrm{d}t} = -\Gamma_{\mathrm{P}}\mathbf{p}(t) + \Gamma_{\mathrm{P}}\mathbf{m}'(t). \end{cases} \tag{16}$$

This is identical to the result in the main paper, since $\mathrm{d}m'/\mathrm{d}t$ can always be evaluated at $t$ rather than at $t+\tau$. In an analogous fashion, the total delay time $\tau$ can be distributed arbitrarily over the transcription and translation process. The resulting phase shift will affect the boundary conditions, and must be taken into account.

### S5. EXPERIMENTAL PARAMETERS

In Stricker *et al.* [9], experiments are presented on a single, negative feedback loop oscillator in the presence of inducer IPTG. We model the inducer as an additional reservoir, the size of which is set by the internal IPTG concentration. Assuming the total number of IPTG molecules is low enough to not signicantly alter the effective concentration of 'free' nonspecific DNA sites, this way of modelling is justified.

The dissociation constant for IPTG to LacI is $5.0 \times 10^{-6}$ M [10–12]. IPTG-bound LacI loses its strong association with operator DNA [13, 14] which indeed induces the redistribution of LacI towards non-specific DNA [15]. The dissociation constant is related to the specific binding free energy with respect to a reference state $\mu^0$ through

$$K_\mathrm{s} = v_\mathrm{w} e^{-\beta(\epsilon_\mathrm{s} - \mu^0)} \tag{17}$$

where $K_\mathrm{s}$ is the reciprocal of $K_\mathrm{D}$. With $v_\mathrm{w} = 18 \times 10^{-3}$ L mol$^{-1}$ this amounts to a binding free energy $\epsilon_\mathrm{IPTG} - \mu^0 = -16.2\ k_\mathrm{B}T$ . It makes sense to set the zero of energy at the effective free binding energy of LacI bound to the non-specific reservoir[16], free of IPTG. As such, we can take the binding free energy of the IPTG competitor site as $\epsilon_\mathrm{IPTG} = -16.2\ k_\mathrm{B}T$.

In the work of Fernández-Castané *et al.* [17], it is stated that at steady-state, the intracellular IPTG concentration in *E. coli* is on the order of 100 µM for an external medium concentration of 0.6 mM IPTG. Taking the cell volume of *E. coli* to be roughly 1.1 µm$^3$[2], then 1.5 nM corresponds to one molecule per cell. This sets the number of IPTG molecules to roughly $5 \times 10^4$.

The supplementary information to Stricker *et al.* [9] reports an experimental doubling period in the order of 30 min, which leads to a protein degradation constant in the order of $(\Gamma_\mathrm{p})_{ii} \approx \log(2)/(30\,\mathrm{min}) \approx 0.023\,\mathrm{min}^{-1}$ . Furthermore, the article reports a mRNA degradation constant of $0.54\,\mathrm{min}^{-1}$. The appropriate time delay was chosen to obtain oscillations with a frequency of $\approx 3\,\mathrm{h}^{-1}$ as per ref [9](supplementary figure 5). We found oscillations with this frequency for a time delay of 6 min.

TABLE S1. Reference values for physical parameters in the experimental oscillator in Stricker *et al.* [9]

| Quantity | Value | Unit | Reference |
|---|---|---|---|
| $\Gamma_{\mathrm{M}}$ | 0.54 | $\mathrm{min}^{-1}$ | Stricker *et al.* [9] |
| $\Gamma_{\mathrm{P}}$ | 0.023 | $\mathrm{min}^{-1}$ | Stricker *et al.* [9] |
| $\tau$ | 6 | min | tuned to Stricker *et al.* [9] |
| $\epsilon_{\mathrm{LacI}}$ | −14.8 | $\mathrm{k_B T}$ | Landman *et al.* [16] |
| $\epsilon_{\mathrm{IPTG}}$ | −16.2 | $\mathrm{k_B T}$ | O'Gorman *et al.* [10], Donner *et al.* [11], Chakerian *et al.* [12] |

In table S1 we list an overview of experimental parameters that were used in the main paper.


[1] F. M. Weinert, R. C. Brewster, M. Rydenfelt, R. Phillips, and W. K. Kegel, Scaling of Gene Expression with Transcription-Factor Fugacity, Physical Review Letters **113**, 258101 (2014).

[2] R. Phillips, J. Kondev, J. Theriot, H. G. Garcia, and N. Orme, *Physical Biology of the Cell*, 2nd ed. (Garland Science, New York, 2012).
